# Discovery of Flaviviridae-derived endogenous viral elements in shrew genomes provide novel insights into *Pestivirus* ancient history

**DOI:** 10.1101/2022.02.11.480044

**Authors:** YQ Li, M Bletsa, Z Zisi, I Boonen, S Gryseels, L Kafetzopoulou, JP Webster, S Catalano, OG Pybus, F Van de Perre, HT Li, YY Li, YC Li, A Abramov, P Lymberakis, P Lemey, S Lequime

**Author notes:** corresponding author: Sebastian Lequime.

## Abstract

As viral genomic imprints in host genomes, endogenous viral elements (EVEs) shed light on the deep evolutionary history of viruses, ancestral host ranges, and ancient viral-host interactions. In addition, they may provide crucial information for calibrating viral evolutionary timescales. In this study, we conducted a comprehensive *in silico* screening of a large dataset of available mammalian genomes for EVEs deriving from members of the viral family *Flaviviridae*, an important group of viruses including well-known human pathogens. We identified two novel pestivirus-like EVEs in the reference genome of the Indochinese shrew (*Crocidura indochinensis*). Homologs of these novel EVEs were subsequently detected *in vivo* by molecular detection and sequencing in 27 shrew species, including 26 species representing a wide distribution within the Crocidurinae subfamily and one in the Soricinae subfamily. Based on this wide distribution, we estimate that the integration event occurred before the last common ancestor of the subfamily, about 10.8 million years ago, attesting to an ancient origin of pestiviruses and *Flaviviridae* in general. Moreover, we provide the first description of *Flaviviridae*-derived EVEs in mammals even though the family encompasses numerous mammal-infecting members, including major human pathogens such as Zika, dengue, or hepatitis C viruses. This also suggests that shrews were past and perhaps also current natural reservoirs of pestiviruses. Taken together, our results expand the current known *Pestivirus* host range and provide novel insight into the ancient evolutionary history of pestiviruses and the *Flaviviridae* family in general.

## 1 Introduction

Endogenous viral elements (EVEs) are integrations of partial or full-length viral genomic material into the host genome (Katzourakis and Gifford, 2010a). In addition to retroviruses, which incorporate their genomic sequences into their host genome as an essential part of their replication cycle, many eukaryotic viruses have been found endogenized in various hosts (Feschotte and Gilbert, 2012). These non-retroviruses can derive from dsDNA (Aswad and Katzourakis, 2014; Li and Li, 2015; Liu et al., 2020), ssDNA (Belyi et al., 2010a; Katzourakis and Gifford, 2010a; Kobayashi et al., 2019), dsRNA (Horie et al., 2010; Katzourakis and Gifford, 2010b; Liu et al., 2012), and even ssRNA viruses (Crochu et al., 2004; Flynn and Moreau, 2019; Horie et al., 2010; Lequime and Lambrechts, 2017). Non-retroviral RNA virus-derived EVEs arise from a conjunction of relatively rare events: i) production of DNA genomic material, using retrotransposon encoded reverse transcriptase (Horie, 2019; Horie et al., 2010), ii) integration in the host chromosome of germ-line cells, and iii) overcoming genetic drift and/or natural selection at the population until fixation (Aiewsakun and Katzourakis, 2015a; Holmes, 2011). EVEs thus reflect long-term and intimate interactions of viruses with their hosts, and their identification can reveal insights into past and present host distributions of viral genera and families. The detection of endogenous bornavirus-like elements in invertebrate genomes (Horie et al., 2013), for example, suggested a broader host range for bornaviruses than previously thought. The study of filovirus-like EVEs in some small mammals offer predictive value for further identifying filovirus reservoirs (Taylor et al., 2010). Similarly, the discovery of flavivirus-derived EVEs in *Anopheles* mosquito genomes supported the idea that *Anopheles* mosquitoes could also be the natural hosts of flaviviruses (Lequime and Lambrechts, 2017), as confirmed by other studies (Colmant et al., 2017; Öncü et al., 2018).

Aside from qualitative insights into host ranges of viruses, EVEs can also shed light on deep evolutionary histories of viruses. EVEs are significant traces of past virus-host interactions; unlike animals or plants, viruses do not leave physical fossil records, limiting our ability to study their deep evolutionary histories. EVEs could thus be considered as “genomic fossil records” that can help to unravel long-term evolutionary dynamics between hosts and viral families. The presence of EVE homologs in different host species hints at an integration event before speciation, here the time to the most recent common ancestor. It thus provides a minimum age estimate for the integration event, and therefore a minimum age for the existence of a specific viral taxonomic group (Aiewsakun and Katzourakis, 2015b). For example, EVEs derived from adeno-associated viruses appear to be orthologous in African and Asian elephants, indicating an integration event more than 6 million years ago (Kobayashi et al., 2019). Similarly, the discovery of abundant bornavirus-like EVEs across vertebrates reveals that ancient bornaviral infections occurred over a timeframe of about 100 million years before present (Kawasaki et al., 2021). These studies can also provide genetic fossil calibration points for further gauging ancient viral timescales using phylogenetics (Feschotte and Gilbert, 2012).

Members of the *Flaviviridae* family are linear, positive-sense, single-stranded RNA viruses currently classified in four recognized genera (*Flavivirus*, *Hepacivirus*, *Pestivirus*, and *Pegivirus*). They encompass many significant pathogens, such as dengue, Zika, hepatitis C, or bovine viral diarrhea virus. In addition to human and livestock infections, *Flaviviridae* viruses have been detected in a broad range of hosts, e.g. non-human primates (Mares-Guia et al., 2020), rodents (Bletsa et al., 2021), bats (Wu et al., 2018), birds (Strand et al., 2018), arthropods (Shi et al., 2016), and fish (Hartlage, 2016; Soto et al., 2020). Divergence date estimates on a genus-wide scale are relatively rare, with the oldest, for the *Flavivirus* genus, being estimated at 200,000 years ago (Pettersson and Fiz-Palacios, 2014). Although *Flaviviridae* viruses have been detected in various hosts, especially in small mammals (Bletsa et al., 2021; de Lamballerie et al., 2002; Wu et al., 2020, 2018), current published studies have only identified *Flaviviridae*-derived EVEs in arthropods, including mosquitos (Crochu et al., 2004; Lequime and Lambrechts, 2017; Roiz et al., 2009), ticks (Maruyama et al., 2014), and crustaceans (Parry and Asgari, 2019), and these are all related to the *Flavivirus* genus. A recent study also identified *Flaviviridae*-derived EVEs in a wide variety of hosts, including various invertebrates and fish (Bamford et al., 2021). Currently, however, convincing evidence for *Flaviviridae*-derived EVEs in vertebrates remains lacking. Interestingly, potential integration has been suggested in medaka fish (Belyi et al., 2010b) as well as in rabbit and hare genomes (Silva et al., 2015, 2012). While this raises the hypothesis that *Flaviviridae* viruses have integrated in vertebrate hosts, the origin of these specific genomic sequences remains inconclusive due to their short size and low sequence similarity to the *Flaviviridae* (*Flavivirus* and *Hepacivirus* respectively).

In this study, we explored *in silico* the presence of *Flaviviridae*-derived EVEs in a comprehensive set of mammalian genomes, and we discovered two novel pesti-like EVEs in the genome of the Indochinese shrew *Crocidura indochinensis.* We subsequently identified and characterized homologs of these EVEs *in vivo* in 26 species of the Crocidurinae subfamily and one member of the Soricinae subfamily, establishing the integration event at least 10.8 million years ago. Our results provide the first evidence for an ancient origin of pestiviruses and also contribute to a better understanding of the evolutionary history of the *Flaviviridae* family in general.

## 2 Material and Methods

### 2.1 *In silico* survey

#### 2.1.1 Data collection

To screen for *Flaviviridae*-like EVEs, 689 mammalian genomes (57 bats, 9 insectivores, 177 rodents, 101 nonhuman primates, 207 even-toed ungulates, 15 odd-toed ungulates, 108 carnivores, and 15 marsupials), were retrieved from the National Center for Biotechnology Information (NCBI) Whole Genome Shotgun (WGS) database (last accessed in November 2020). A detailed list of all the surveyed mammalian genomes is provided in Supplementary Table S1. A representative group of 306 *Flaviviridae* or *Flaviviridae*-like polyprotein sequences was compiled from the NCBI non-redundant protein database (accessed in February 2019). We provide a list of the nucleotide/protein accession numbers in Supplementary Table S2.

#### 2.1.2 Genome screening

*Flaviviridae* polyprotein sequences were used as queries in tBLASTn (BLAST+ v2.6.0) (Camacho et al., 2009) searches with mammalian genomes as targets. To avoid potential artifacts, only hits with E-value < 10^−4^ and length >= 250 nt were extracted from mammalian genomes based on the reported position by BLAST in the host contig. These putative EVEs were then used as query in a reciprocal tBLASTx (BLAST+ v2.6.0) (Camacho et al., 2009) against a local NCBI nucleotide (nt) database (accessed in October 2018) and BLASTx (BLAST+ v2.6.0) (Camacho et al., 2009) against a non-redundant protein (nr) database (accessed in October 2018). EVEs were confirmed if the best hits contained *Flaviviridae* family members with an E-value < 10^−4^ and length >= 250 nt. The presence of conserved viral genetic features within the hits was assessed using the NCBI Conserved Domain Database (Marchler-Bauer et al., 2015).

#### 2.1.3 EVE characterization

Upon identification of the EVEs, they were translated and aligned with corresponding polyprotein sequences from several representative *Flaviviridae* species using MAFFT v7.453 (Katoh et al., 2002). All alignments were trimmed in BMGE v1.12 (Criscuolo and Gribaldo, 2010) in order to select for phylogenetic informative regions. The best substitution models were PMB+G4 for the EVE1 alignment, LG+F+G4 for the EVE2 alignment, and LG+F+I+G4 for the concatenated EVEs alignment according to the BIC criterion and were used to construct phylogenetic ML trees with IQ-TREE v1.6.12 (Nguyen et al., 2015).

#### 2.1.4 Flanking region analysis

To characterize the EVEs loci and identify potential transposable elements or other genetic features, flanking regions of the identified EVEs were extracted from the host contigs and used as BLAST queries to screen against the NCBI nucleotide (nt) and non-redundant protein (nr) databases (both accessed in October 2018).

#### 2.1.5 Metagenomic screening

According to the WGS screening results above, some *Flaviviridae*-related hits were detected in a shrew (*Crocidura indochinensis*) genome. However, apart from the *Crocidura indochinensis* and *Sorex araneus* complete genomes, only a limited number of shrew genomes are currently available in the NCBI WGS database. Therefore, 73 DNA experimental genomic data sets and 6 RNA-Seq transcriptome data sets (Supplementary Table S3) from the Soricidae family were retrieved from NCBI Sequence Read Archive (SRA) database using SRA Toolkit v2.10.8 (Leinonen et al., 2011). Reads were mapped to the identified EVEs nucleotide references using Bowtie2 v2.3.5.1 (Langmead and Salzberg, 2012). Alignment files were processed with SAMtools v1.10 (Li et al., 2009) and coverage was determined using bedtools v2.27.1 (Quinlan and Hall, 2010) and visualized in RStudio v1.1.463.

### 2.2 *In vivo* validation

#### 2.2.1 Sample collection

Based on the screening results, to further verify the presence of *Flaviviridae*-related EVEs *in vivo*, a total of 65 tissue and DNA samples from species belonging to the *Crocidura* genus and 6 other related genera of the Soricidae family, namely *Paracrocidura*, *Scutisorex*, *Suncus*, *Sylvisorex* (subfamily Crocidurinae), *Neomys,* and *Sorex* (subfamily Soricinae), were screened for the presence of the identified EVEs. These samples were previously collected in China, Vietnam, Africa, and the Eastern Mediterranean (Supplementary Table S4) as part of other studies (Bannikova et al., 2011; de Perre et al., 2019; Jenkins et al., 2013; van de Perre et al., 2018)

#### 2.2.2 Target EVEs and cytochrome b (Cytb) amplification

DNA was extracted from tissue samples using the DNeasy Blood & Tissue Kit (Qiagen) following the manufacturer’s instructions.

To screen for the presence of EVEs *in vivo*, we designed 18 PCR primers (Supplementary Table S5) spanning the 2 EVEs region and a section of the intermediate flanking region from the host genome. Amplicons were generated with DreamTaq DNA Polymerase (ThermoFisher Scientific) using the following cycling conditions: (i) 3 min of denaturation at 95°C, (ii) 35 cycles of 95°C for 30s, 56°C for 30s, 72°C extension for 1min/kb, and (iii) 10 min final extension at 72°C.

To confirm the host species, and to complement available specimen information, the mitochondrial Cytb gene was amplified using general primers (Supplementary Table S5) of the Crocidurinae subfamily. PCR reactions were conducted using the DreamTaq DNA Polymerase (ThermoFisher Scientific) with the following thermal cycling conditions: (i) 3 min of denaturation at 95°C, (ii) 35 cycles of 95°C for 30s, 56°C for 30s, 72°C extension for 1min/kb, and (iii) 10 min final extension at 72°C.

All PCR products were purified using the ExoSAP-IT PCR Product Cleanup (ThermoFisher Scientific) or Zymoclean Gel DNA Recovery Kits (ZYMO Research) to remove primer dimers and unspecific products, following the manufacturer’s instructions.

#### 2.2.3 Sanger sequencing

The generated PCR products were sequenced by Macrogen Europe. The amplicons were mapped to the whole EVEs region (2,235nt) from the WGS *Crocidura indochinensis* contig, and concatenated based on consensus sequence to get the complete EVEs in Geneious Prime® v2020.2.4. Cytb amplicons (~1,140nt) were forward and reverse sequenced and a consensus sequence was generated using Geneious Prime® v2020.2.4.

#### 2.2.4 MinION sequencing

For 12 samples with relatively low-quality Sanger sequencing chromatograms (additional information provided in Supplementary Table S6), MinION sequencing was performed to obtain the complete EVEs region (~2,235nt) together with the Cytb gene (~1,140nt). The Oxford Nanopore Technologies (ONT) 1D Native barcoding genomic DNA protocol was used without the DNA fragmentation step and the barcoded amplicons were loaded onto the MinION device. We used the MinKNOW software v19.13.5 on the MinIT companion for data acquisition and basecalling. Qcat v1.1.0 (ONT, https://github.com/nanoporetech/qcat) was used to demultiplex reads under the epi2me algorithm and to trim bad quality reads and adapters with min score of 90. The EVE regions extracted from the WGS *Crocidura indochinensis* contig were used as references to map the reads with Minimap2 v.2.22 (Li, 2018) using -ax map-ont parameters. Alignments were converted and indexed using SAMtools v1.10 (Li et al., 2009) and consensus sequences were generated using a custom Python script (Kafetzopoulou, 2019).

### 2.3 Phylogenetic analysis and visualization

All generated EVEs sequences were translated and aligned with homologous polyproteins from available *Pestivirus* species using MAFFT v7.453 (Katoh et al., 2002). Sequences of dengue-2 and Zika virus (*Flavivirus* genus) were used as an outgroup. The alignment was trimmed using BMGE v1.12 (Criscuolo and Gribaldo, 2010) and the filtered regions were used to construct maximum-likelihood (ML) phylogenetic trees using IQ-TREE v1.6.12 (Nguyen et al., 2015) under the best-fitting models (according to the Bayesian information criterion): WAG+G4 (for EVE1), LG+F+I+G4 (for EVE2), and LG+G4 (for complete concatenated EVEs). Phylogenies were visualized and annotated using FigTree v1.4.4 (A. Rambaut; http://tree.bio.ed.ac.uk/software/figtree/). Percent identity matrices were generated using Clustal Omega (Sievers et al., 2011) via EMBL-EBI web services (Madeira et al., 2019).

Cytb sequences of EVEs-positive specimens (n=48) were aligned using MAFFT v7.453 (Katoh et al., 2002) together with a dataset of *n*=393 Soricidae nucleotide sequences downloaded from NCBI. The generated alignment was trimmed in BMGE v1.12 (Criscuolo and Gribaldo, 2010) and an ML phylogeny was reconstructed using IQ-TREE v1.6.12 (Nguyen et al., 2015) with the best-fitting model (TIM2+F+I+G4).

To highlight the evolutionary relationships of our newly discovered EVEs and their hosts, EVEs and Cytb phylogenies were annotated in ggtree v1.14.6 (Yu et al., 2017) and treeio v1.6.2 (Wang et al., 2020) R packages, followed by the estimation of a co-phylogenetic plot (tanglegram) using the ape v5.0 (Paradis and Schliep, 2019) and dendextend v1.14.0 (Galili, 2015) R packages.

### 2.4 Characterization of selective pressure

We only characterized the selective pressure acting on the EVE 1 locus, as the EVE 2 locus exhibits widespread stop codons and translation frame shifts. We aligned the open reading frame of the complete EVE1 region (318 nt) using MEGA v11.0.9 (Tamura et al., 2021). An ML tree was build based on this alignment in IQ-TREE v1.6.12 (Nguyen et al., 2015) under the best-fitting model (HKY+F). We then conducted two site-specific selection analyses to characterize the selective pressure on each site using estimates of the ratio of non-synonymous/synonymous substitution rate (*ω* = *d*_*N*_/*d*_*S*_): 1) fixed effects likelihood (FEL) analyses (Kosakovsky Pond and Frost, 2005) using MG94xREV model available in HyPhy software v2.5.3 (Kosakovsky Pond et al., 2005), and 2) the Bayesian renaissance counting method (Lemey et al., 2012) implemented in BEAST v1.10.5 (Suchard et al., 2018). The value of *ω* quantifies the selective pressure, with *ω* > 1 suggesting positive selection, *ω* = 1 neutral evolution and *ω* < 1 negative or purifying selection.

## 3 Results

### 3.1 False positive Flaviviridae-like hits from mammalian genome screening

Our initial screening of 689 available mammalian genomes using *Flaviviridae* and *Flaviviridae*-like polyproteins yielded 66 positive hits from 49 species, including rodents, non-human primates, marsupials, insectivores, carnivores, and bats. Detailed information about our *in silico* screening results can be found in Supplementary Table S7. All hits were similar to three pestiviruses, namely border disease virus, bovine viral diarrhea virus 1, and bovine viral diarrhea virus 2. With the exception of one shrew species (see section below), the position of all hits in the corresponding viral genomic sequence was in the ubiquitin-homolog domain between the nonstructural proteins NS2 and NS3, while some of the hits slightly expanded the alignment to the NS3 region (shown in Supplementary Fig. 1). The ubiquitin domain in bovine viral diarrhea virus is however predicted to originate from cellular derived insertions in cytopathogenic pestivirus (Agapov et al., 2004; Becher and Tautz, 2011). The similarity between this viral genomic region and ubiquitin poses a considerable risk for false positives when searching for pestivirus-derived EVEs, and these hits were therefore not further considered.

### 3.2 *In silico* identification of *Crocidura indochinensis* pesti-like EVEs

Besides the ubiquitin-related false positive results, our *in silico* screening identified a series of five *Flaviviridae*-related EVEs fragments in a single contig of the *Crocidura indochinensis* reference genome PVKC01 (Table 1). The first EVE (EVE1) is 318 nt long, with its closest BLAST hit being the Linda virus (Pestivirus) envelope glycoprotein E2 region (tBLASTx, 25.5% identity, e-value 3.31E^−35^), without any stop codon (Fig. 1). The remaining four EVEs fragments are 1,053 nt, 84 nt, 114 nt and 87 nt long respectively, with their closest BLAST hits being a classical swine fever virus and a rodent pestivirus (with minimum amino-acid identity 24.9%, maximum 62.1%). These four fragments are separated by very short gaps, with lengths of 16, 1 and 4 nucleotides, respectively. The two central fragments of EVE2, fragments 3 (84 nt) and fragment 4 (114 nt), are in a different translation frame than the two others, but in the same orientation. Their arrangement in the host contig reflects their relative position in the pestiviral genome, which partially spans the non-structural NS2 and NS3 genes (Fig 1). For these reasons, in our further analyses, we considered these four fragments as the result of a single pestiviral integration event, and thus a unique EVE (EVE2). Phylogenetic reconstructions of the identified *Crocidura indochinensis* EVE1, EVE2 and entire concatenated sequence with exogenous *Flaviviridae* viruses supports the pestivirus-origin of the EVEs (Supplementary Fig. 2).

**Table 1.**
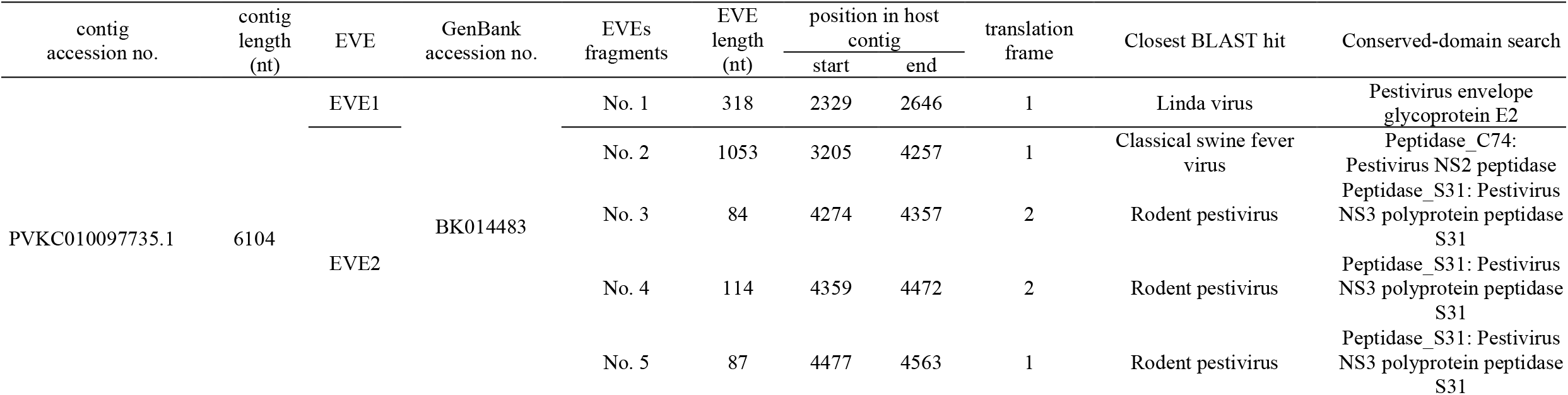
Newly detected EVEs in *Crocidura indochinensis* genome.

**Fig. 1.**
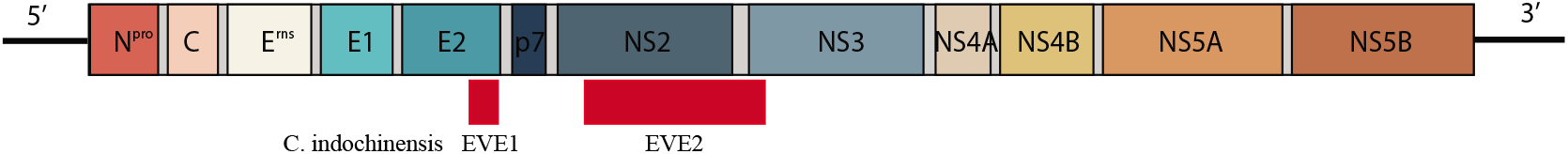
The positions of the newly detected *Crocidura indochinensis* EVEs are shown relative to an archetypal *Pestivirus* genome (classical swine fever virus, NC_002657). N^pro^, N-terminal protease; C, nucleocapsid core protein; E^rns^, envelope glycoprotein E^rns^; E1, envelope glycoprotein E1; E2, envelope glycoprotein E2; p7, nonstructural protein p7; NS2, nonstructural protein NS2; NS3, nonstructural protein NS3; NS4A, nonstructural protein NS4A; NS4B, nonstructural protein NS4B; NS5A, nonstructural protein NS5A; NS5B, nonstructural protein NS5B.

In addition, a fragment prior to EVE1 in the contig shows a strong similarity (tBLASTx, 45.2% identity, e-value 5.51E^−19^, supplementary Table 7) with the pestivirus ribonuclease T2 gene. However, considering that this enzyme exists in a wide range of organisms (Luhtala and Parker, 2010), the virus-derived origin of this sequence in *Crocidura indochinensis* is not guaranteed. The position of all pestivirus-like hits, including the ribonuclease T2 gene, in the host contig corresponds to their relative organization in the pestivirus genome. No additional features were detected after a tBLASTx search of the whole contig encompassing the two identified EVEs (Supplementary Table S7). In addition, we did not detect the EVE sequences in reads of publicly available experimental genomic and transcriptomic data from Soricidae species.

### 3.3 Identification and distribution of pesti-like EVEs in other Soricidae species

To expand our screening and evaluate the distribution of these newly identified EVEs in additional species that are phylogenetically close to *Crocidura indochinensis*, we undertook a PCR-based screening of 65 samples from 29 species of the Soricidae family (Supplementary Table S4). These samples belonged to 7 different genera (*Crocidura*, *Paracrocidura*, *Scutisorex*, *Suncus*, *Sylvisorex*, *Neomys,* and *Sorex*), encompassing two subfamilies, Crocidurinae and Soricinae. Cytb genomic sequences were also generated to confirm the species identification (Supplementary Table S6 & Supplementary Fig. 3).

In total, 58 samples derived from 27 species contained the newly identified pesti-like EVEs, and 48 samples yielded complete or nearly complete EVEs sequences, representing 22 species in the Crocidurinae subfamily and one species (*Neomys anomalus*) in the Soricinae subfamily (Fig. 2). All novel EVEs sequences were highly similar to the *Crocidura indochinensis* EVEs sequences, with a mean identity of 90.38% on amino acid level, and 95.12% on nucleotide level, respectively (Supplementary Table S8). Phylogenetic reconstruction indicated that all Crocidurinae pesti-like EVEs clustered together as a sister-lineage of currently recognized *Pestivirus* species (Fig. 3, Supplementary Fig. 4).

**Fig. 2.**
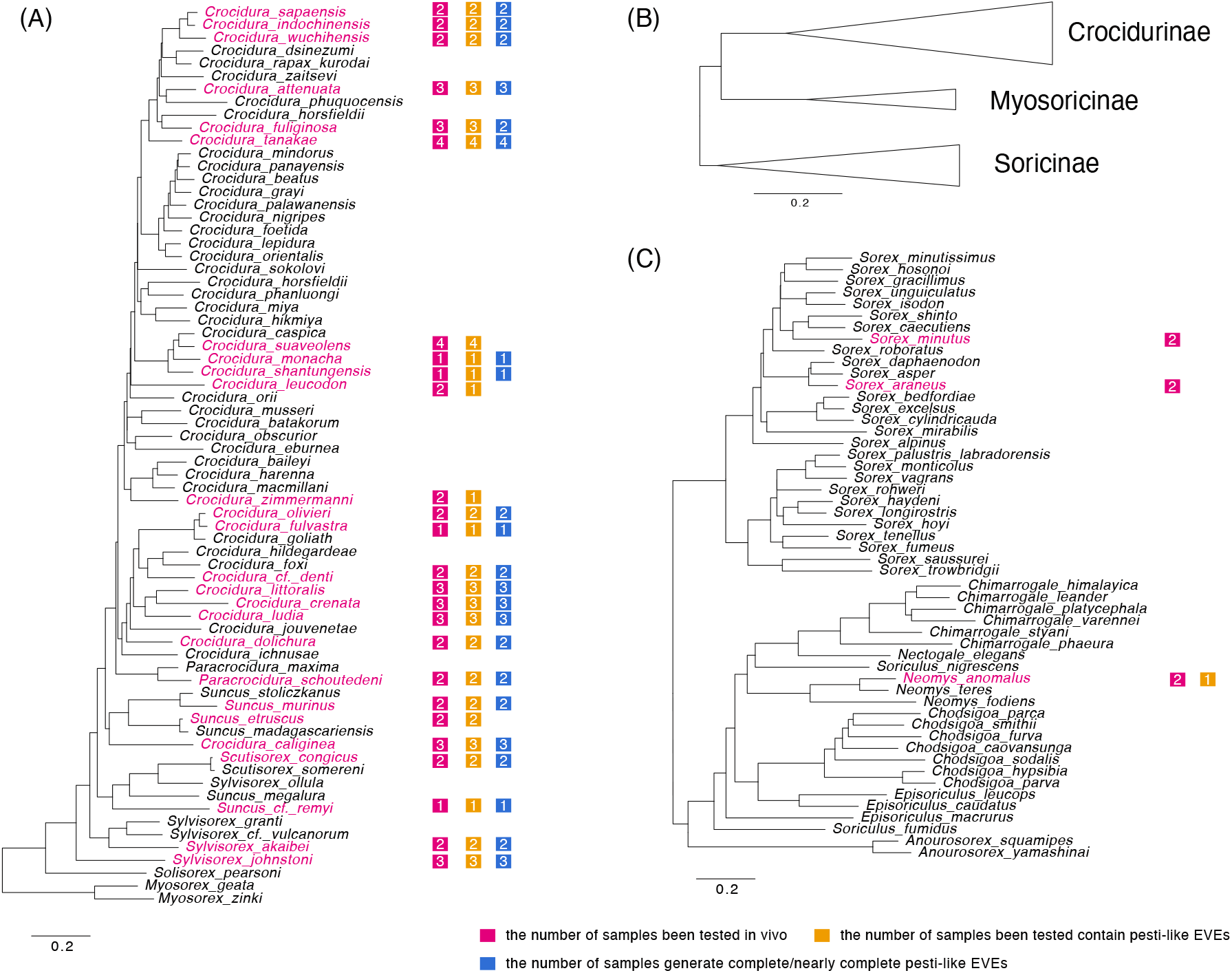
Maximum likelihood phylogeny of the Soricidae (shrews) family based on the Cytb gene and the available samples’ species distribution in this study, highlighted in pink: (A) The phylogeny of Crocidurinae subfamily and sample distribution; (B) Subfamilies relationships within the Soricidae family; (C) The phylogeny of Soricinae subfamily and sample distribution.

**Fig. 3.**
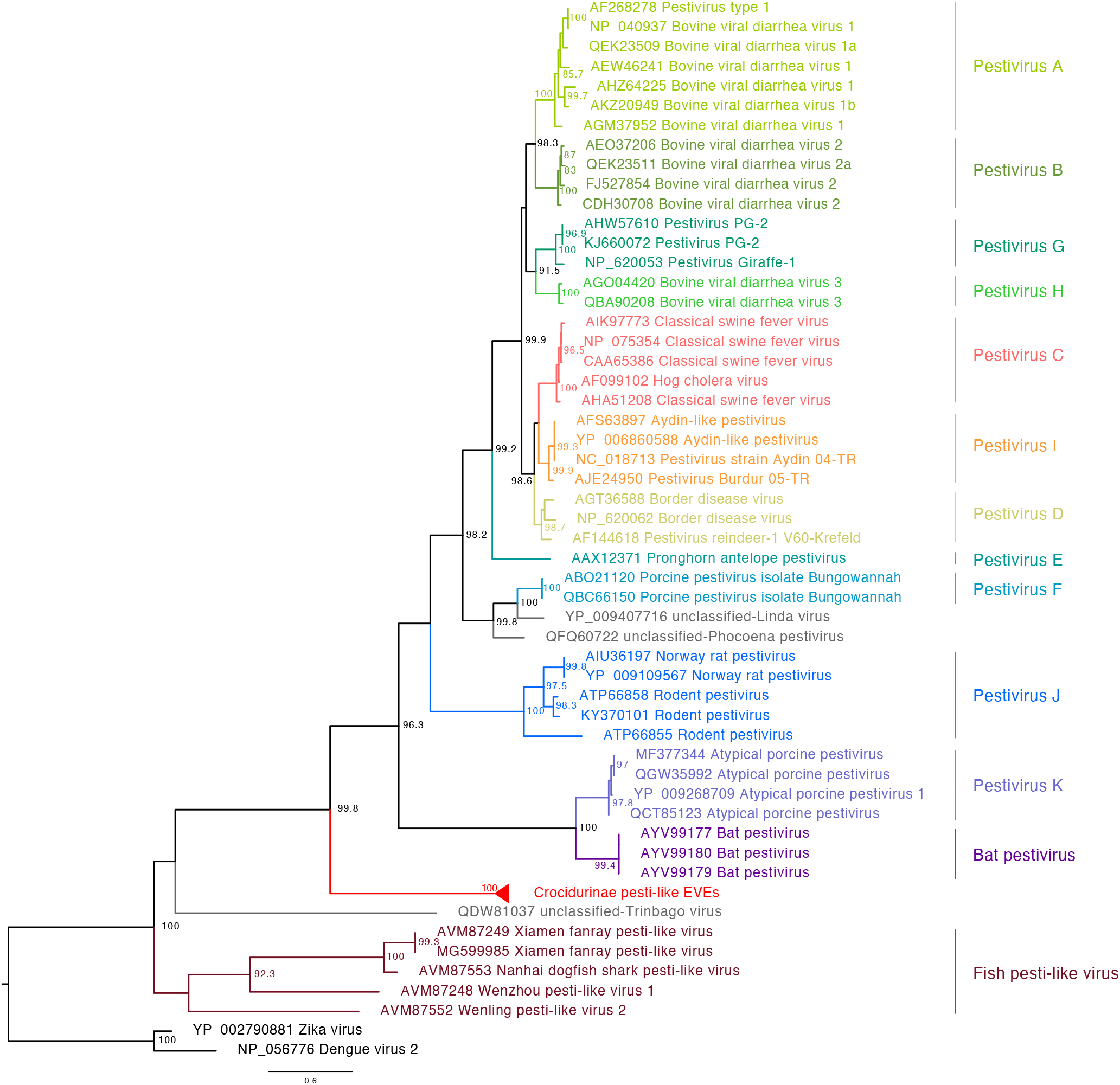
Phylogenetic relationships of pesti-like EVEs with representative *Pestivirus* species and with Dengue and Zika virus (*Flavivirus*) as outgroup. Clades are colored based on viral species. Node labels indicate Shimodaira-Hasegawa (SH)-like branch support (%, only values > 80% are shown). Scale bars indicate the number of amino acid substitutions per site.

Though collected in different locations, nearly all species tested in the Crocidurinae subfamily harbored the pesti-like EVEs sequences. For some species however, such as *Crocidura* cf. *zimmermanni*, *C. leucodon*, *C. suaveolens* and *Suncus etruscus*, we could not always detect or sequence the EVEs in all samples. The failed detection could be explained by the genomic template being of poor quality due to storage conditions associated with the museum specimens. Interestingly, the pesti-like EVEs were also detected in one *Neomys anomalus* sample, while the remaining Soricinae specimens yielded negative results. The widespread nature of these homolog EVEs in the Crocidurinae species suggests a single endogenization event before their common ancestor about 10.8 million years ago (Dubey et al., 2007). Since we did not manage to sequence the pesti-like EVEs found in *Neomys anomalus*, their phylogenetic relationship to the Crocidurinae pesti-like EVEs is unclear and may not necessarily derive from the same endogenization event. To compare the evolutionary history between the EVEs and the Cytb gene, we constructed a tanglegram based on their respective phylogenies, which illustrates specific phylogenetic discordances that may also be expected between mitochondrial and nuclear genes (Fig 4).

**Fig. 4.**
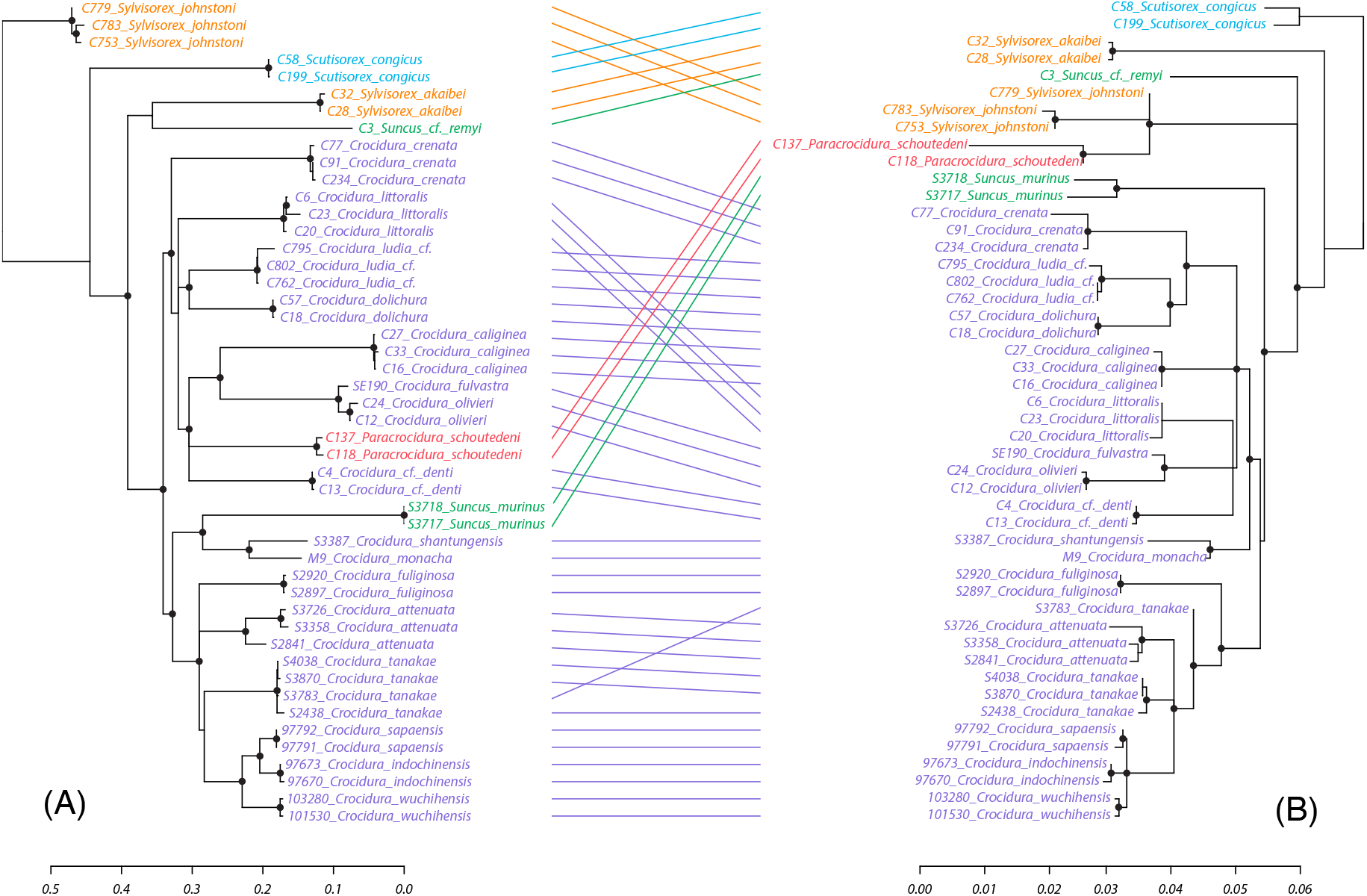
Tanglegram of the cytochrome b phylogeny (A) and the corresponding EVEs phylogeny (B). The cytochrome b tree was inferred for gene sequences from 22 shrew species, and the EVEs tree (right) was inferred using the newly generated EVEs sequences. The clade nodes with SH-like branch support < 50% were collapsed as polytomies. The black circles indicate the SH-like branch support > 80%. Lines connect corresponding tips in the two phylogenies. Scale bars indicate the number of nucleotide substitutions per site.

### 3.4 Selective pressure on pesti-like EVE1 locus

For testing the indication of function of the EVEs region, we further processed selective pressure analysis. We only characterized the selective pressure acting on the EVE1 locus, as the EVE2 locus exhibits widespread stop codons and translation frame shifts. We measured the selective pressure acting on EVE1 by estimating the ratio (*ω*) of non-synonymous substitution rate (*d*_*N*_) and synonymous substitutions rate (*d*_*S*_) in protein-coding sequences using two methods: 1) fixed effects likelihood (FEL) (Kosakovsky Pond and Frost, 2005), and 2) Bayesian renaissance counting (Lemey et al., 2012). It is expected that non-functional region should conform to neutral evolution while functional region would appear to be under purifying selection. Based on the 48 pesti-like EVE1 potentially coding sequences the FEL method indicates an overall neutral evolution with *ω* = 0.94. Likewise, the Bayesian renaissance counting model yields a ratio at 1.098 [95% credible interval: 0.658, 1.624], reflecting neutral evolution.

## 4 Discussion

Non-retroviral endogenous viral elements (EVEs) are rare traces of the ancient evolutionary history of viruses. These genomic fossils offer valuable insights into host range, ancestral genetic diversity and can provide invaluable information for dating viral evolutionary history (Aiewsakun and Katzourakis, 2015a; Feschotte and Gilbert, 2012). In our study we screened a comprehensive set of mammalian genomes to discover such *Flaviviridae*-derived EVEs. We uncovered two *Flaviviridae*-derived EVEs sequences in the genome of the Indochinese shrew and confirmed their presence in a broad range of shrew species belonging to the Crocidurinae subfamily.

The EVEs we identified are related to extant viruses within the *Pestivirus* genus. Viruses belonging to this genus were initially detected in a variety of artiodactylous hosts, such as ruminants and swine, in which they cause subclinical or clinical infections including hemorrhagic syndrome, abortion, acute fatal mucosal disease. Recent metagenomic studies extended the host range towards rodents (Wu et al., 2020, 2018), bats (Wu et al., 2018), fish (Shi et al., 2018), and ticks (Sameroff et al., 2019), but to some extent, the restricted sampling beyond agriculturally important animals limits our understanding of the real host range. Shrews, for example, have been recently identified as host of hepaciviruses, another genus in the *Flaviviridae* family (Guo et al., 2019; Wu et al., 2020), but to date not of pestiviruses. The broad detection of pestivirus-derived EVEs reported in our study strongly supports that ancestors of the Crocidurinae shrew subfamily have been hosts of pestiviruses and suggests that their descendants might still be. Indeed, considering the extremely low probability of a non-retroviral endogenization event to occur in the germline, EVEs are strong indicators of frequent interactions between the original exogenous viruses and their hosts (Aiewsakun and Katzourakis, 2015a; Feschotte and Gilbert, 2012). Although direct detection and characterization of pestiviruses from shrews are still required to formally demonstrate that they are natural hosts of pestiviruses, our study provides indirect support for a wider and more diverse host range of pestiviruses.

Given the low probability of endogenization events of non-retroviral RNA viruses and the contiguous nature of the two EVEs on the host and viral genome, our results suggest a single endogenization event followed by genetic drift. One or several insertion events separated the original EVE in two fragments, EVE1 and EVE2. EVE1 shows a short but intact open reading frame to be evolving under neutral evolution while EVE2 exhibits multiple stop codons and frame-shifts due to additional insertions. Many studies have identified the important roles that EVEs can play in host antiviral immunity, both in vertebrates and invertebrates (Blair et al., 2020; Ophinni et al., 2019; Skirmuntt et al., 2020). Flavivirus-like EVEs in *Aedes* mosquitoes, for example, can produce P-element-induced wimpy testis (PIWI)-interacting RNAs (piRNAs) which limit the cognate virus replication (Suzuki et al., 2020). It is highly unlikely that the pesti-like sequences we discovered have a function in shrews because of the absence of negative selection and the disruption of the original viral coding region.

The evolutionary history of the EVEs sequences after integration remains unclear. Interestingly, they do not show any appreciable patterns of phylogenetic consistency with the host Cytb gene sequences. The EVE sequences might not be a reliable genetic marker to discriminate species (Tobe et al., 2010), as highlighted by the low amount of genetic diversity compared to Cytb sequences (Fig 4). Additionally, the discrepancy might be explained by differences in the genetic inheritance of the Cytb gene and EVEs: the Cytb gene is a mitochondrial gene whereas EVEs are integrated in the nuclear genome. This can lead to different observed evolutionary patterns, especially in the case of weak reproductive isolation within species or species complex, allowing hybridization, as has been suggested for some *Crocidura* species (Dubey et al., 2008, 2006; Vogel et al., 2004).

Dating the ancient evolutionary history of ssRNA viruses such as pestiviruses and *Flaviviridae* in general is challenging. The most commonly used method for inferring viral divergence time is based on the estimation of evolutionary rates derived from sequence data and their collection dates. However, the applicability of this method is often limited by heterogeneous substitution rates though time (Aiewsakun and Katzourakis, 2016) and among viral lineages (Duffy et al., 2008; Sanjuán, 2012). Not accounting for the former leads to recent estimates for the origins of ssRNA viruses that are often in conflict with other phylogenetic evidence (Holmes, 2003). Using suitable molecular clock models, the powerful combination of both tip and node calibrations may help to recover more accurate evolutionary timescales (O’Reilly et al., 2015). Node calibration is however challenging for viruses as no fossil evidence can be found. It thus often relies on known phylogeographic events and other indirect calibrations point, such as ecological events or assumptions of co-divergence as alternative (Bamford et al., 2021; Moureau et al., 2015; Pettersson and Fiz-Palacios, 2014). The discovery of ssRNA virus-related EVEs thus enable a direct estimation for a robust long-term timeline of virus evolution history by co-phyletic analysis of EVE’s orthologs in different host (Gilbert and Feschotte, 2010).

The pesti-like EVE sequences characterized in our study are widespread in Crocidurinae species, are monophyletic and exhibit high sequence similarity. Considering the low probability of endogenization events of non-retroviral RNA viruses, this suggests that the pesti-like EVE got integrated before the most-recent common ancestor of the subfamily, which is estimated to be over 10.8 million years ago (Dubey et al., 2007). There are only a handful of molecular dating estimates for pestiviruses and they mostly focus on viral species or clades that are associated with economic losses. Diversification of bovine viral diarrhea virus 1 (Pestivirus A) subtypes was estimated to have started about 363 years ago (Weber et al., 2021), and the divergence of HoBi-like pestivirus (Pestivirus H) was dated back to the 16^th^ century (Silveira et al., 2020). Our results provide the first robust ancient time node for pestiviruses based on the estimated EVEs integration date and suggest that pestiviruses were already circulating in mammals more than 10.8 million years ago.

In conclusion, we discovered and characterized the first *Flaviviridae*-related EVEs records from mammalian reference genomes, which derived from pestiviruses. The wide EVEs distribution in shrew Crocidurinae subfamily indicates they are a historical host group of pestiviruses and further suggests a robust ancient origin time of the *Pestivirus* genus. Our results show the key role of EVEs not only in expanding our knowledge about ancient viral-host interactions, but also their importance in reconstructing the viral evolutionary history, which contributes to our understanding of viral evolutionary dynamics from ancient times to the present.

## 5 Acknowledgement

The samples from Vietnam were collected during biodiversity surveys carried by the Joint Vietnam-Russian Tropical Research and Technological Centre.

## 6 Funding

The research leading to these results has received funding from the European Research Council under the European Union’s Horizon 2020 research and innovation programme (grant agreement no. 725422-ReservoirDOCS). PL acknowledges support by the Research Foundation -- Flanders (`Fonds voor Wetenschappelijk Onderzoek -- Vlaanderen’, G066215N, G0D5117N and G0B9317N). YQ LI acknowledges the support of China Scholarship Council. AVA acknowledges the support by Ministry of Science and Higher Education of the Russian Federation (grant no. 075-15-2021-1069). YC LI acknowledges the support of National Natural Science Foundation of China (grant ID. 31672254). F.V.d.P. acknowledges the support of the Ph.D. fellowship from the Research Foundation– Flanders. M.R.W.

## Notes

### Competing Interest Statement

The authors have declared no competing interest.

### Summary of Updates

Updated author list and affiliation

