## Supplementary Fig. 1 for "Discovery of Flaviviridae-derived endogenous viral elements in shrew genomes provide novel insights into *Pestivirus* ancient history"

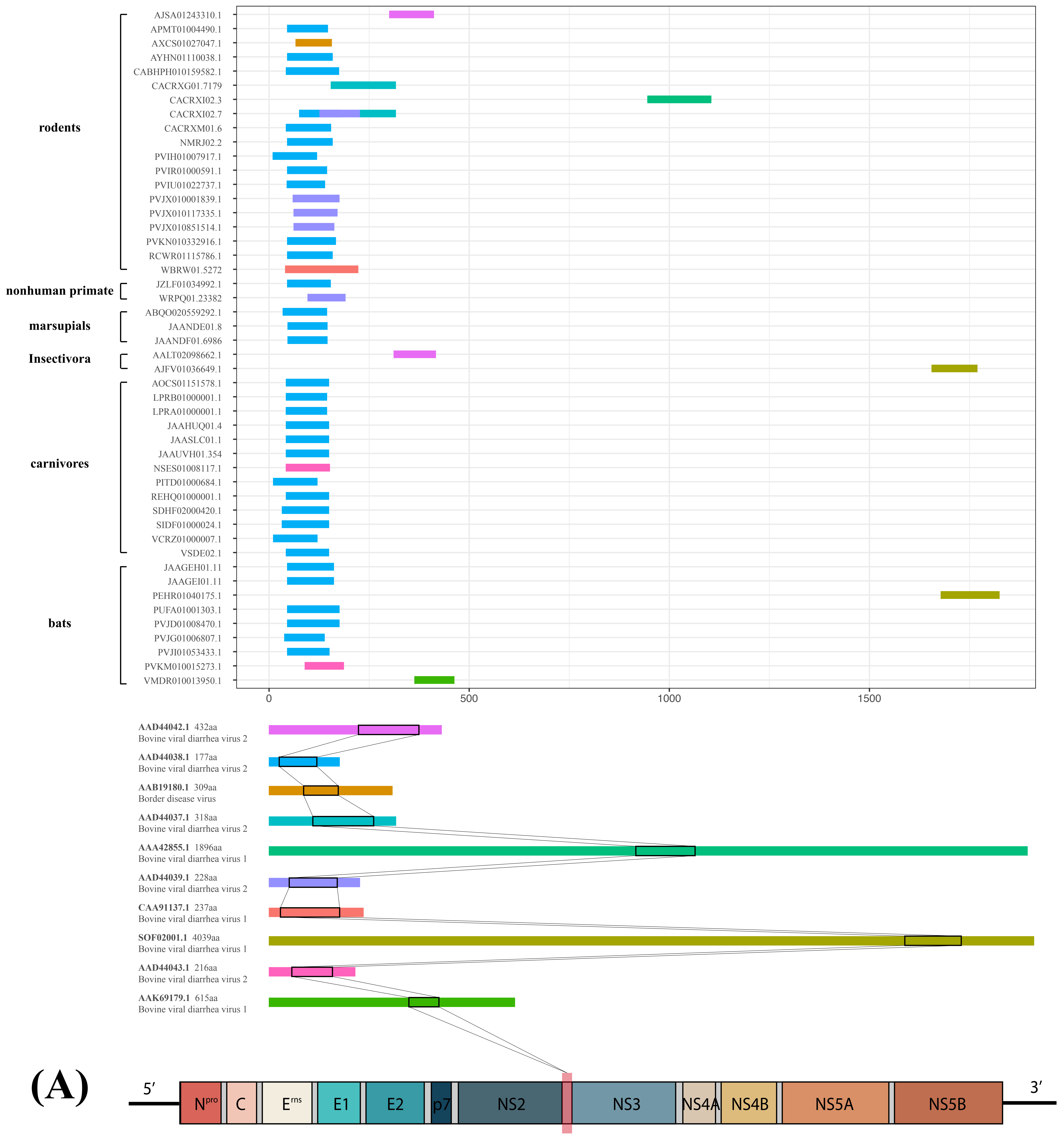


**Supplementary Fig.1: (A) Positive hits result from screening against NCBI nr database and the corresponding position in specific pestiviral genome**

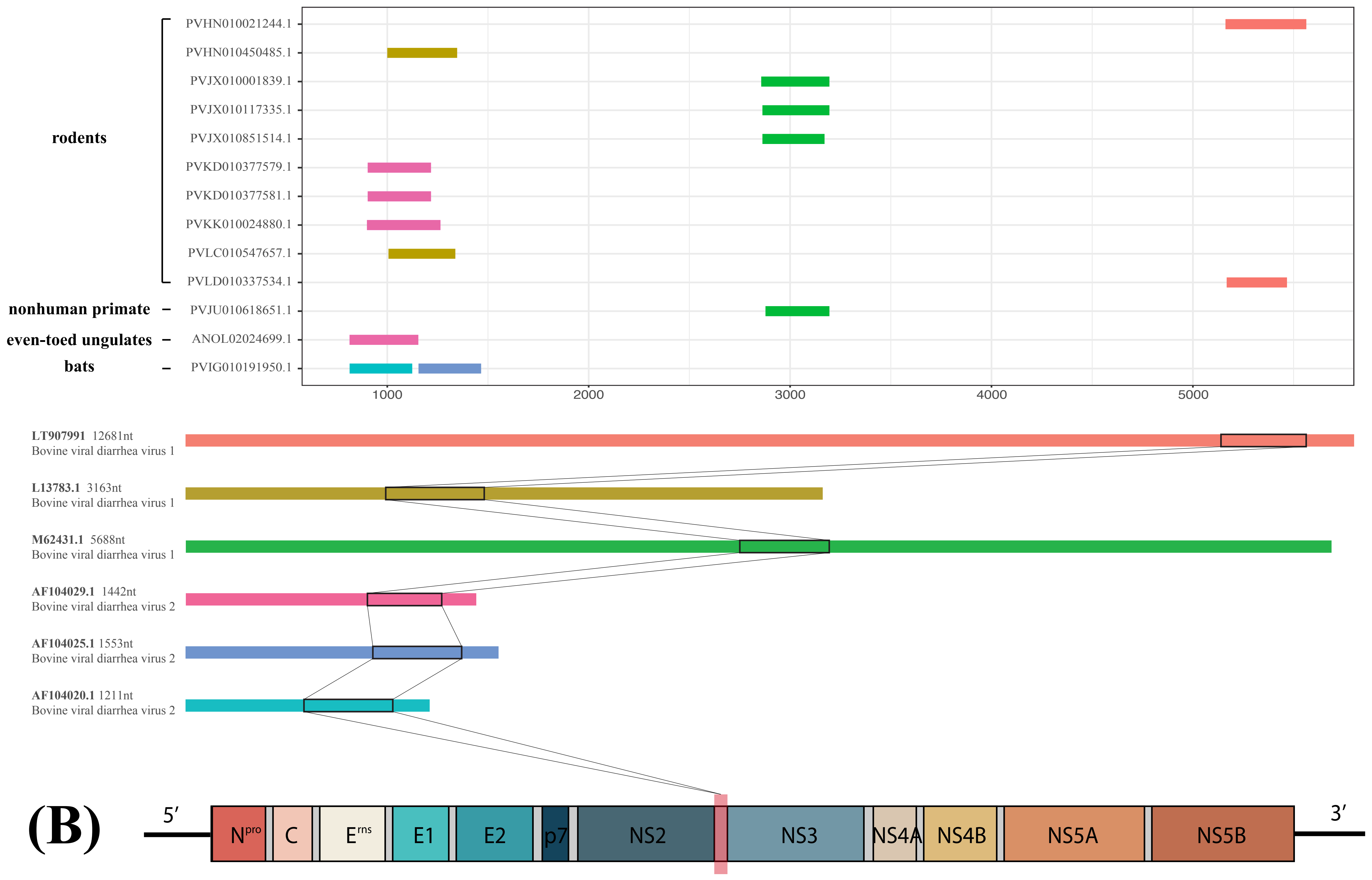


**Supplementary Fig.1: (B) Positive hits result from screening against NCBI nt database and the corresponding position in specific pestiviral genome**
