## Supplementary figures and images for "Discovery of Flaviviridae-derived endogenous viral elements in shrew genomes provide novel insights into *Pestivirus* ancient history"

### Supplementary Fig. 2

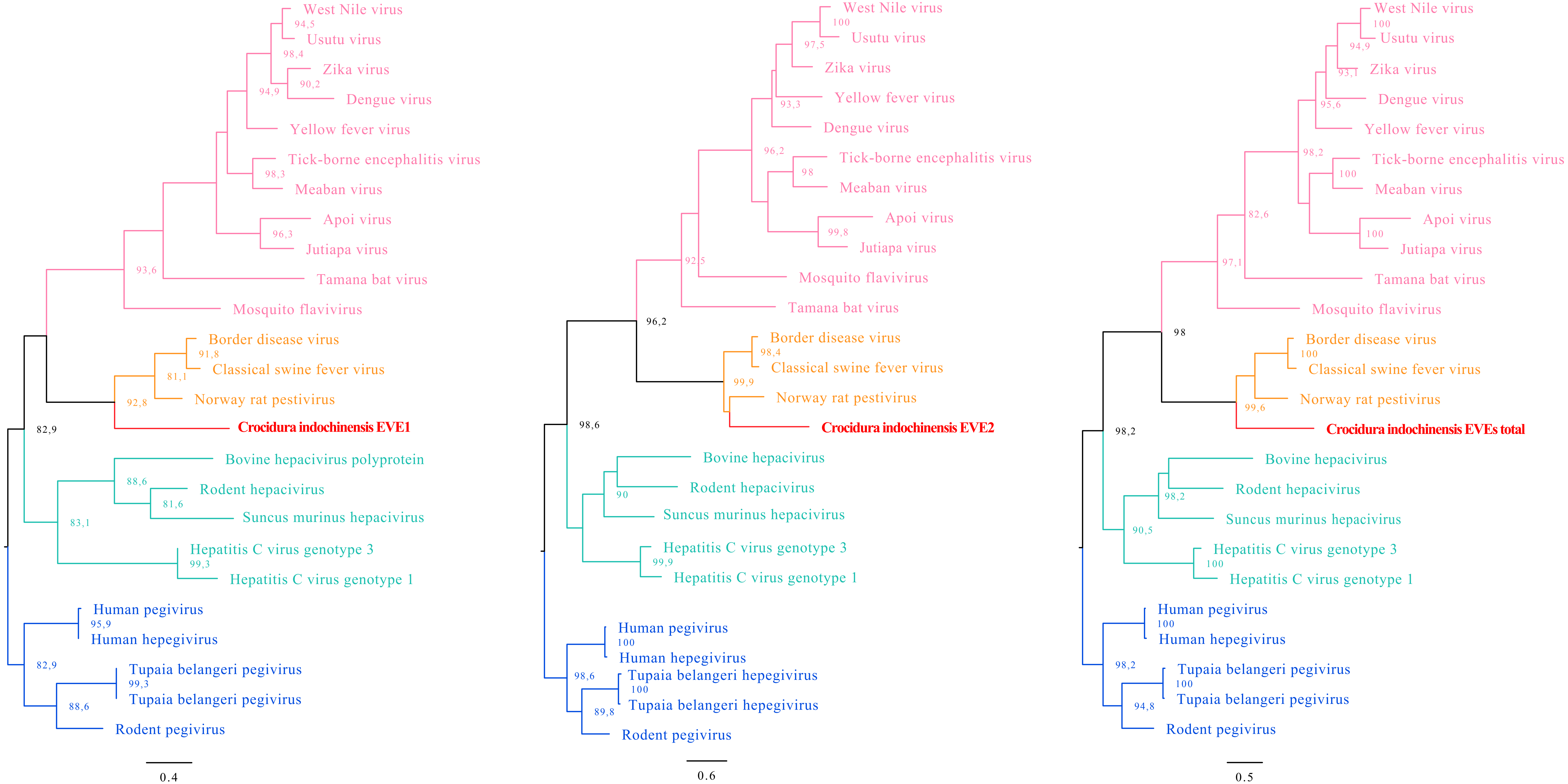

### Supplementary Fig. 3

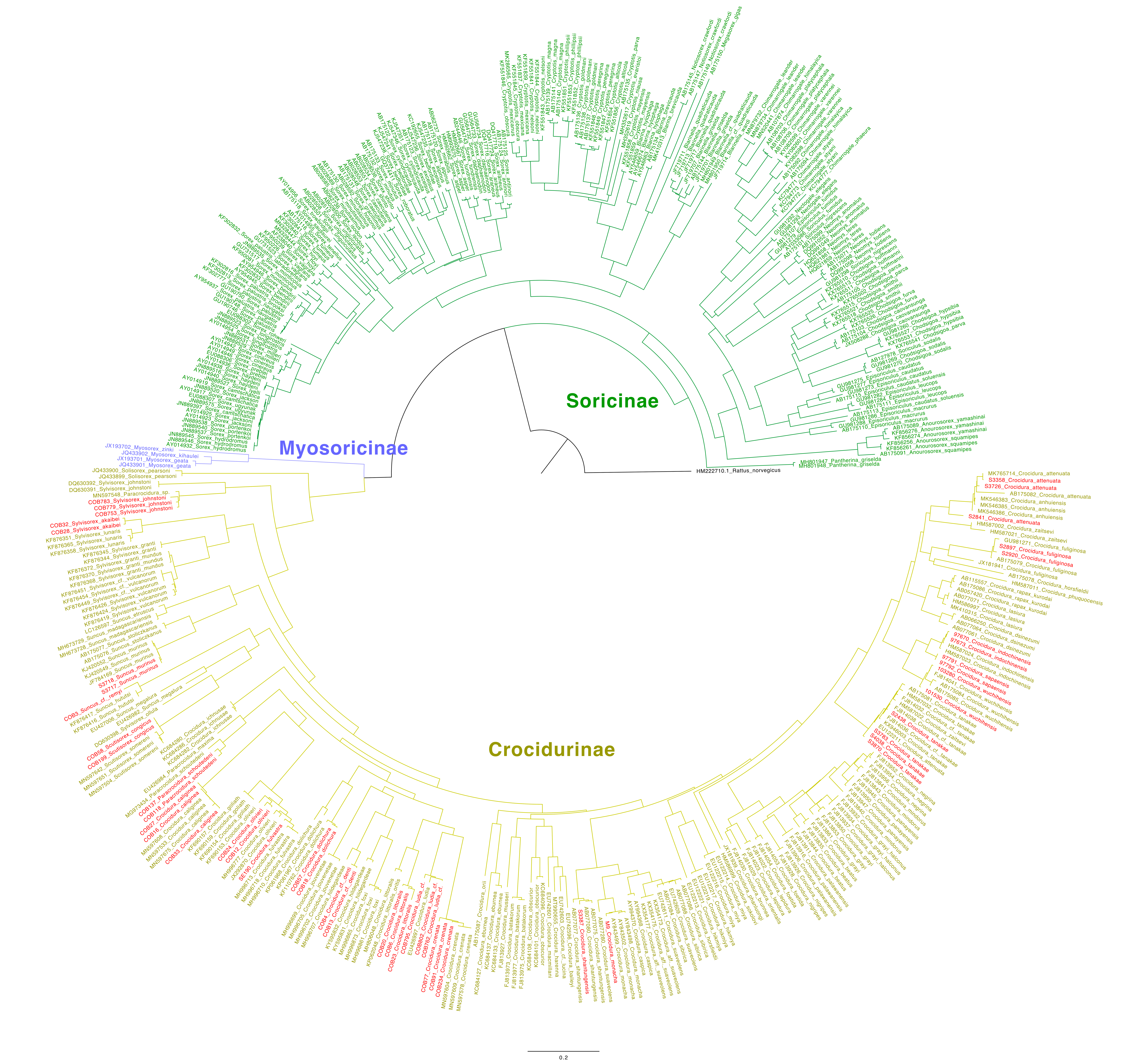
