## Supplementary Fig. 4 for "Discovery of Flaviviridae-derived endogenous viral elements in shrew genomes provide novel insights into *Pestivirus* ancient history"

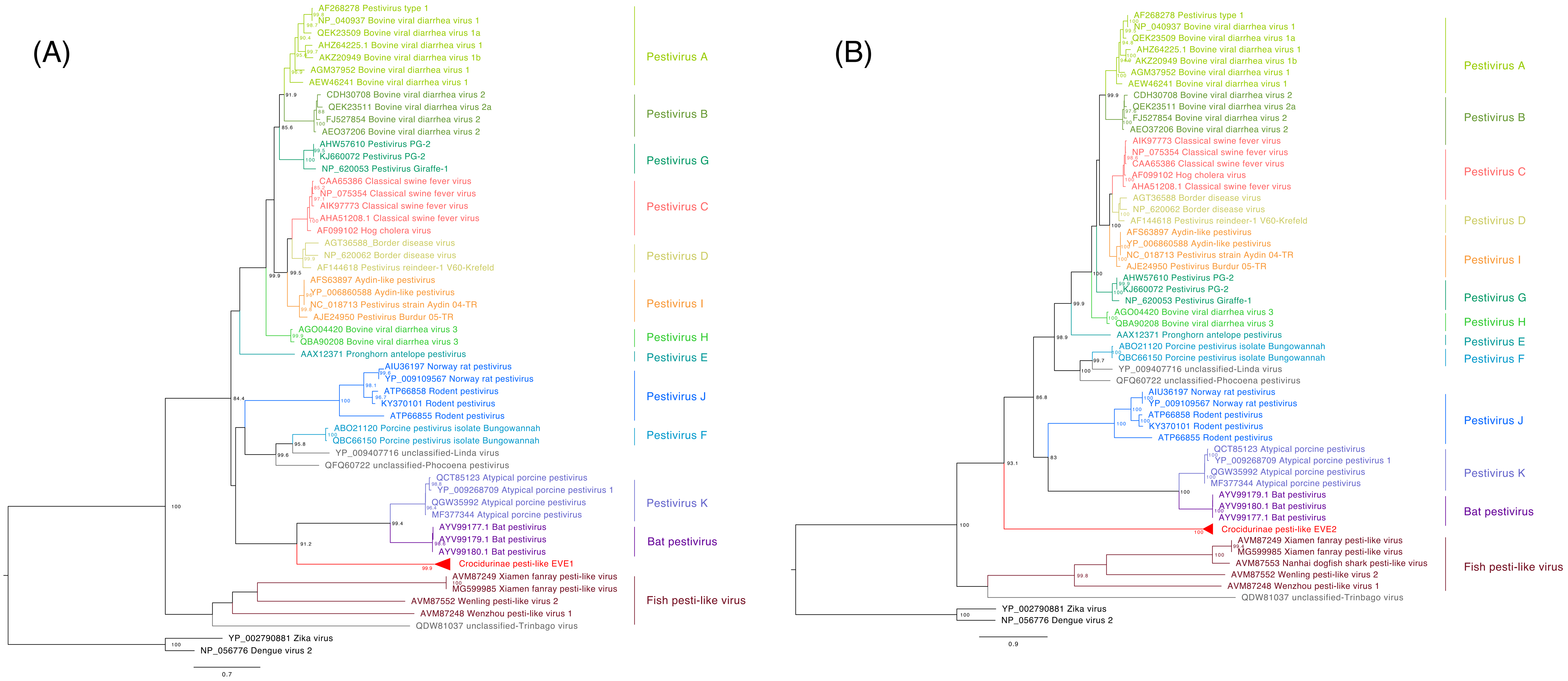


**Supplementary Fig. 3: Phylogenetic relationships of pesti-like EVE1 region (A) and EVE2 region (B) with representative Pestiviruses, with Dengue and Zika virus (Flavivirus) as outgroup. Clades are colored based on viral species. Node labels indicate Shimodaira-Hasegawa (SH)-like branch support (%, only values > 80% are shown). Scale bars indicate the number of substitutions.**
